# NON-TARGETED METABOLOMICS REVEAL DIFFERENCES IN THE METABOLIC PROFILE OF THE FALL ARMYWORM STRAINS WHEN FEEDING DIFFERENT FOOD SOURCES

**DOI:** 10.1101/2022.05.18.492515

**Authors:** Nathalia C. Oliveira, Larry Phelan, Carlos A. Labate, Fernando L. Cônsoli

**Author notes:** **Correspondence** Fernando L. Cônsoli, Insect Interactions Laboratory, Department of Entomology and Acarology, Luiz de Queiroz College of Agriculture, University of São Paulo, Piracicaba, São Paulo, Brazil.

## Abstract

*Spodoptera frugiperda*, the fall armyworm (FAW), is an important polyphagous agricultural pest feeding on nearly 350 host plants. FAW is undergoing incipient speciation with two well-characterized host-adapted strains, the “corn” (*CS*) and “rice” (*RS*) strains, which are morphologically identical but carry several genes under positive selection for host adaptation. We used non-targeted metabolomics based on gas chromatography/mass spectrometry to identify differences in metabolite profiles of the larval gut of *CS* and *RS* feeding on different host plants. Larvae were fed on artificial diet, maize, rice, or cotton leaves from eclosion to the sixth instar, when they had their midgut dissected for the analysis. This study revealed that the midgut metabolome of FAW varied due to larval diet and differed between the FAW host-adapted strains. Additionally, we identified several candidate metabolites that may be involved in the adaptation of *CS* and *RS* to their host plants. Our findings provide clues toward the gut metabolic activities of the FAW strains.

## 1. INTRODUCTION

*Spodoptera frugiperda*, the fall armyworm (FAW) (Lepidoptera, Noctuidae), feeds on approximately 350 different species from 76 families (Montezano et al., 2018). Despite this wide range of host plants, FAW is best known as one of the most important agricultural pests of grasses (maize, millet, rice and sorghum) and some cultivated dicots such as cotton (Barros et al., 2010). The FAW is native to the New World, but in the last few years has invaded Africa and further spread to Asia and Oceania (Goergen et al., 2016; Johnson, 1987; Otim et al., 2018; Piggott et al., 2021; Suby et al., 2020), Therefore, FAW is currently considered of a global concern due its polyphagy and capacity for rapid evolution of resistance to pesticides and Bt crops (Huang, 2021; Jakka et al., 2016), representing an imminent threat to food security and a source of significant economic losses.

The FAW is the only species of *Spodoptera* that usually feeds on grasses without having adapted, suitable mandibles. Larvae that feed on grasses typically have specialized mandibles with chisel-like edges adapted to the consumption of silica-rich leaves, which cause wear to larval mandibles(Djamin and Pathak, 1967; Pogue, 2002; Smith, 2005). Mandibles of the *FAW* have serrate-like processes adapted to the consumption of dicots or monocots that do not accumulate silica (Pogue, 2002). FAW is primitively polyphagous, but because of the mandible-type it is thought to have started exploiting cultivated grasses as host plants only recently (Kergoat et al., 2021, 2012).

Another interesting aspect of the FAW is the identification of two distinct strains known as the rice (*RS*) and corn (*CS*) strains (Gouin et al., 2017; Pashley, 1986). There are indications that this divergence occurred about 2 Myr ago (Kergoat et al., 2021, 2012).These strains differ in their performance and preference for host plants, and the correct classification of these two strains of the FAW is still controversial. Some authors refer to them as “sibling species” (Drès and Mallet, 2002; Dumas et al., 2015), “host strains” (Pashley, 1986; Prowell et al., 2004), “host form” (Juárez et al., 2014) and “morphocryptic strains” (Sarr et al., 2021).

The lack of consensus is due to the fact these strains co-exist in sympatry and still hybridize, but also due to inconsistencies in the associations with the named host plants. At the adult stage, both corn and rice strains showed weak evidence of preference for their expected host plant in choice and non-choice laboratory experiments (Orsucci et al. 2020). Despite the fact that the corn strain is often associated with maize, sorghum, and cotton, with the rice strain associated with rice and pasture grasses, some reports show the rice strain larvae developed better on corn and sorghum than corn strain larvae (Meagher et al. 2004). Moreover, both strains performed poorly when feeding on rice (Silva-Brandão et al. 2017). Therefore, further studies are still needed to understand how the process of host-plant adaptation is taking place in FAW.

Every novel acquisition of host plant by herbivores constitutes a new niche adaptation program that opens several evolutionary possibilities, but not without associated costs. In order to exploit a novel host, insects have to become adapted to deal with new defensive secondary metabolites, such as phenolics and terpenoids, and the nutritional quality of the new host plant (Singer, 2008). However, the mechanisms behind the best performance of a given host-adapted strain on a given plant are poorly understood so far. Different approaches can be used to address this question. One alternative is to access the insect metabolome, the set of all low-molecular-weight metabolites that are produced during cell metabolism (Sun and Hu, 2016). Ultimately, the metabolome is a product of genomic, transcriptomic, and/or proteomic processes (Johnson and Gonzalez 2012). Non-targeted metabolomics provides a holistic view of the insect metabolic profile. It makes no assumptions about which metabolites are important in distinguishing sample types (Sévin et al., 2015). This approach provides a direct functional measurement of cellular activity and physiological state, reflecting environmental changes such as new host plants as well as aspects related to their genome, as different host-adapted strains. Therefore, the non-targeted study of metabolomes is a good tool to highlight candidate metabolites involved in insect-plant interactions (Maag et al., 2015). Particularly, the assessment of the insect midgut may be useful, bearing in mind that it is a selectively permeable and metabolically active tissue, in which most digestion and almost all nutrient absorption takes place (Dow, 1987). However, approaches focused on the assessment of the gut metabolomics of insect herbivores are not common, and little is known on how host plants impact the metabolic profile of the herbivore gut.

The gut microbiome is also a key player in th e metabolic processes of their hosts. Gut microbes can play important roles in several metabolic functions, including vitamin production (Chen et al., 2016; Salem et al., 2014), amino acid synthesis (Ayayee et al., 2016; Xia et al., 2017) and detoxification of secondary plant compounds and synthetic insecticides (Almeida et al., 2017; Ceja-Navarro et al., 2015). Among the numerous factors that influence the gut microbiota (R J Dillon and Dillon, 2004; Yun et al., 2014), diet has received considerable attention due to its strong effect on the composition of the microbial community (Mason et al., 2020; Wongsiri and Randolph, 1962; Yun et al., 2014). Diet provides the substrates to produce a plethora of small molecules that can be converted by the gut microbiota, and which are not produced by the host (Krishnan et al., 2015; Wang et al., 2020). Therefore, the gut microbiota may also facilitate adaptation to new host plants by regulating or participating in the host’s metabolic processes (Hammer and Bowers, 2015; Zhang et al., 2020). Microbial contribution will depend on substrate availability and on microbial gene diversity and activity (Wu et al., 2016). Thus, taxonomic or metagenomic information of the gut microbiota is limited in predicting the metabolome of a microbial community, as it may under- or overestimate the functional contribution of associated gut microbiota depending on the nutritional conditions the host is exposed to (Wu et al., 2016).

The FAW is a good model to study adaptation of phytophagous insects to agricultural plants. Moreover, the metabolic processes underlying host shifts or differentiation in this species are not well understood. In terms of metabolome, we would expect different metabolic profiles to reflect new adaptations. We predict differences in larval adaptation to host plants to be reflected in the metabolome of their gut. These differences might highlight adaptations in response to plant chemistry, changes in metabolic pathways, and/or new roles for microbial symbionts. The aim of the present research is to investigate if the gut metabolome of FAW is determined by the diet and/or by host genotype. Highlighting the metabolic differences in the midgut of the FAW strains may represent the starting point for future research that aims to clarify : 1) how different host plants affect insect nutritional metabolism and 2) how larvae of the two host strains differ in their metabolism of different host plant chemistries.

## 2. MATERIAL AND METHODS

### 2.1 Insect rearing and strains identification

Colonies of FAW were initiated in the laboratory from field-collected populations. The *RS* was originally obtained from rice fields in Santa Maria, RS, Brazil (29°68’68”S, 53°81’49”W) and the *CS* from a maize field in Piracicaba, SP, Brazil (22°43’30”S, 47°38’56”O). Field-collected larvae were individualized into plastic cups containing an artificial diet based on wheat germ, beans and brewer’s yeast (Burton and Perkins, 1972; Kasten Jr et al., 1978), brought to the laboratory and reared under controlled conditions (25 ± 1^?^C; 70 ± 10% RH; 14 h photophase) until pupation. Pupae were transferred to clean plastic cups lined with filter paper for adult emergence. The produced exuviae were used for DNA extraction for strain identification as described below. After strain identification, all newly emerged adults belonging to the same strain were transferred to PVC tubes lined with paper as a substrate for egg laying. Egg masses were collected and transferred to artificial diet for later larval development.

Strain identification followed Levy et al. (2002) (Levy et al., 2002). DNA was extracted from individual pupal exuviae using the genomic DNA preparation protocol from RNAlater™ preserved tissues with some modifications. The exuviae were individually placed in 750 μL digestion buffer (60 mM Tris pH 8.0, 100 mM EDTA, 0.5% SDS) containing proteinase K at a final concentration of 500 μg/mL. Samples were macerated using plastic pestles and mixed well by inversion. Samples were incubated overnight at 55°C. Afterwards, 750 μL of phenol: chloroform (1:1) was added and samples were rapidly inverted for 2 min before centrifugation at a tabletop centrifuge at maximum speed (10 min). The aqueous layer was recovered and subjected to re-extraction twice before a final extraction with chloroform. The aqueous layer was collected and added to 0.1 volume of 3 M sodium acetate (pH 5.2) and an equal volume of 95% ethanol. Samples were then mixed by inversion, incubated for 40 min at −80°C, and centrifuged (27,238 g x 30 min x 4°C). The pellet obtained was washed twice with 1 mL of 85% ice-cold ethanol, centrifuged for 10 min after each wash and dried at 60°C during 5-10 min in a SpeedVac. Finally, the pellet obtained was resuspended in nuclease-free water. DNA concentration and quality were estimated by spectrophotometry and agarose gel electrophoresis (Sambrook and Russell, 2001).

Polymerase chain reaction (PCR) amplification of the mitochondrial COI gene was conducted using the primers set JM76 (5’-GAGCTGAATTAGGRACTCCAGG-3’) and JM77 (5’- ATCACCTCCWCCTGCAGGATC-3’) to produce an amplicon of 569 base pairs (bp)(Levy et al. 2002). The PCR mixture contained 100-150 ng of gDNA, 1.5 mM of MgCl_2_, 1 x PCR buffer, 0.2 mM of each dNTP, 0.32 µM of each primer and 0.5U of GoTaq® DNA Polymerase (Promega) in a total volume of 25 µL. The thermocycling conditions were one cycle at 94°C x 1 min followed by 33 cycles at 92°C x 45 s, 56°C x 45 s, 72°C x 1 min, with a final extension at 72°C x 3 min (1x). Amplicons were then subjected to endonuclease restriction analysis using *Msp*I (HpaII) to produce two fragments (497pb and 72pb) for amplicons of the *CS*, while no digestion is observed for *RS* amplicons. After the amplification, 10 μL of the PCR reaction mixture was subjected to digestion with 10 U of *Msp*I following the manufacture guidelines (product number ER0541®, Thermo Scientific). Samples were gently mixed, centrifuged for a few seconds and incubated overnight at 37°C. Subsequently, digestion efficiency and the resulting products were verified using a 1.5% agarose gel electrophoresis.

### 2.2 Assay with natural diet and gut dissection

Maize (*Zea mays*, family: Poaceae) var. “Conventional impact” and cotton (*Gossypium hirsutum*, family: Malvaceae) var. IAC FC2 seeds were seeded in 500 mL plastic pots filled with soil conditioner, while rice seeds (*Oriza sativa*, family: Poaceae) var. BRS Esmeralda were seeded in 1 L plastic pots. All plants were maintained in a greenhouse.

The leaves were cut and immersed in a container with distilled water for 30 minutes to maintain turgidity. Newly emerged larvae of *RS* and *CS* strains were placed in 25 mL plastic cups containing a 3-cm piece of the host plant leaf (cotton and maize - leaves from v3-v4 stages; rice - v11-13). The leaves were replaced with fresh ones according to the needs of each instar and were replaced every other day or earlier to avoid food shortages. Insects were kept under controlled laboratory conditions throughout the experiments (25 ± 1°C; 60 ± 10%; 14-hour photophase).

The experimental groups were represented by larvae of rice (*RS*) and corn (*CS*) strains reared on the following substrates: rice (RiRS; RiCS), corn (CoRS; CoCS), cotton (CtRS; CtCS), and artificial diet (DiRS; DiCS). We used five replicates for each treatment, with each replicate corresponding to a pool of midgut collected from five larvae.

The guts were collected from sixth instars larvae, one day after the molting. Larvae were surface-sterilized in cooled 0.2% sodium hypochlorite in 70% ethanol and washed in cold sterile water. Surface-sterilized larvae were dissected in sterile water under aseptic conditions. Gut tissues with the peritrophic matrix and the enclosed intestinal content were flash-frozen in liquid nitrogen and stored at −80 until metabolite extraction.

### 2.3 Metabolite extraction

Samples were subjected to metabolite extraction and analysis according to (Hoffman et al., 2010), with some modifications. A pool of midguts from five larvae were macerated in liquid nitrogen, and 25 mg of the macerate was homogenized in TissueLyser II (QIAGEN) at the highest speed for 1 min using 5 mm tungsten beads in 500 μL of methanol-chloroform-water solution (3:1:1). Then the sample was sonicated (60 Hz.s^-1^ x 30 min) in an ice bath (4ºC) and centrifuged (16,000 g x 10 min x 4ºC). The supernatant was collected and filtered through a Luer Lock 0.22 μM filter (Millex®, JBR6 103 03) directly into amber glass vials.

Aliquots (50 μL) of each sample were freeze-dried in a Terrone model LS 3000 lyophilizer, and subjected to derivatization with 30 μL methoxyamine-HCl (20 mg.mL^-1^) in pyridine for 16 h at room temperature. Trimethylsilylation was accomplished with the addition of 1% trimethylchlorosilane (TMCS) in 30 μL of *n*-methyl-*n*-(trimethylsilyl)-trifluoroacetamide (MSTFA) to the samples, followed by incubation for 1 h at room temperature. After silylation, 30 μL of heptane was added to samples, which were immediately analyzed in a random order in a 7890A Agilent Gas Chromatograph coupled to a Pegasus HT TOF Mass Spectrometer (LECO, St. Joseph, MI, USA) (GC-TOF/MS) (Technologies). Samples were injected together with mix of *n*-alkanes standards (C_12_ - C_40_) for the correct calculation of the retention times. Derivatized samples (1 μL) were injected in splitless mode using an automatic sampler-CTC Combi Pal Xt Duo (CTC Analytics AG, Switzerland) coupled to the GC-MS system equipped with two silica columns in line. The first was a DB 5 column (20 m long x 0.18 mm internal diameter x 0.18 μm thick) (Agilent J & W Scientific, Folsom, CA, USA), and the second a Rxi-17 column (0.84 m long x 0.1 mm internal diameter x 0.1 μm thick) (Restek Corporation, Bellefonte, PA, USA). The injector was set at 280°C, the septum bleed rate was 20 mL.min^-1^ and began after 250s from the start of data acquisition (Budzinski et al., 2019), The gas flow was 1 mL.min^-1^. The temperature of the first column was maintained at 80°C for 2 min and increased at 15°C.min^-1^ to 305°C, with a 10 min hold. The temperature of the second column was maintained at 85°C for 2 min, and then raised to 310°C at 15°C.min^-1^, with a 10 min hold. The column effluent was introduced into the ionization source of the Pegasus HT TOF MS. The transfer line and ionization source temperatures were held at 280 and 250°C, respectively. The ions were generated by an electron source (70-eV) at an ionization current of 2.0 mA, and 20 spectra.s^-1^ were acquired in a mass range of 45-800 *m/z*, with the detector voltage set to 1500 V.

### 2.4 GC-TOF/MS data processing

The processing of GC-TOF/MS data was performed in two steps. Initially the generated chromatograms were exported to the ChromaTOF program, version 4.32 Software (LECO, St. Joseph, MI, USA), in which base line correction, deconvolution of the spectra, retention rate correction (RI), retention time correction (RT), peak identification, and alignment and identification of metabolites were processed using the NIST library, version 11. Only metabolites with three or more characteristic masses and a score of 700 or higher were considered valid. Isomers were manually checked and merged. Feature intensities were normalized to total ion chromatogram (TIC).

MetaboAnalyst 5.0 was used to perform all the downstream analyses (Chong et al., 2019). Data was *log*-transformed and scaled using Pareto. Hierarchical clustering was also performed with the *hclust* function in package stat (R v3.5.1) using Ward as a clustering algorithm and Euclidean distances as measures. Sample clustering was presented as a dendrogram. Principal Component Analysis (PCA) was performed in order to separate and classify the sample groups. Type 1 two-way ANOVA was used to examine the effects of strain and diet, and their interaction on metabolite abundance (Xia et al., 2011). False discovery rate was applied to adjust the *p*-values (0.05). Heatmap was built based only on the significant features from ANOVA. The distance measure used was the Euclidean, and Ward was used for clustering algorithm. In order to identify the features that were potentially significant in discriminating the strains on each host plant, pairwise analysis was performed using volcano plot, which combine fold change (FC ≥ |2.0|) and *t*-test analysis, that can control the false discovery rate (*p* ≤ 0.05).

## 3. RESULTS

The host plants on which FAW larvae fed significantly affected the midgut metabolome. The midgut metabolomes of *RS* and *CS* larvae also differed when feeding on the same diet. However, the food source had a greater impact shaping the gut metabolome of FAW than the host strain (show stats on which this is based, e.g., 2-way ANOVA of summed or overall effects). Metabolomic analyses led to the identification of two major clusters of metabolites, allowing the clear separation of larvae fed on artificial diet when compared to those fed on natural diets (Fig. 1A). The metabolomic profile obtained clearly separated FAW larvae fed on monocots (corn and rice) from those feds on the dicot cotton. Additionally, the profile of metabolites obtained for each FAW strain on each food source also led to their clear separation (Fig.1A). The PCA analysis explained a small percentage of the variation among the samples. The first two principal components explained 39.5% of the variation in the data and showed the clustering of the samples according to food source, but the separation based on strains was not as evident as shown in the dendrogram.

**Figure 1:**
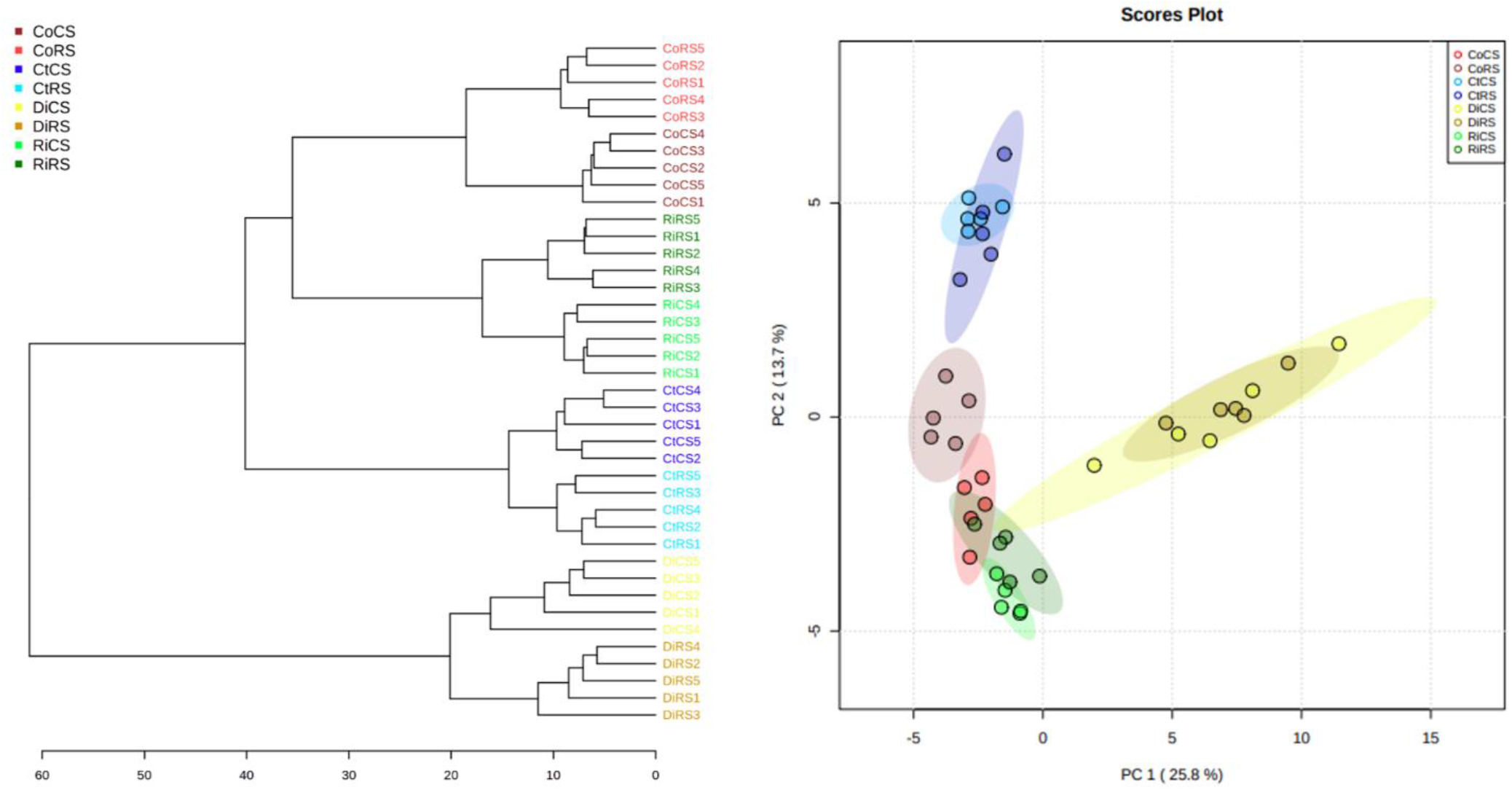
Clustering result shown as dendrogram with distance measure using Euclidean and clustering algorithm using ward.D of the midgut metabolite profiles of *S. frugiperda* larvae (A). Principal Component Analysis (PCA) Scores plot between the selected PCs. The explained variances are shown in brackets (B). The experimental groups were rice strain on rice (RiRS), corn (CoRS), cotton (CtRS) and artificial diet (DiRS) and corn strain on the same diets (RiCS, CoCS, CtCS and DiCS).

Among the 340 peaks identified in the gut samples of FAW larvae, 122 metabolites passed the filter criteria for analysis, with the abundance of 107 them being affected by the diet, 13 by the host-adapted strain, and 50 by the interaction of both factors (Table 1). The abundance of 12 metabolites was simultaneously affected by diet, strain, and their interactions (Fig. 2). The compounds were predominantly classified as amino acids, sugars, fatty acids, and organic acids.

**Table 1.**
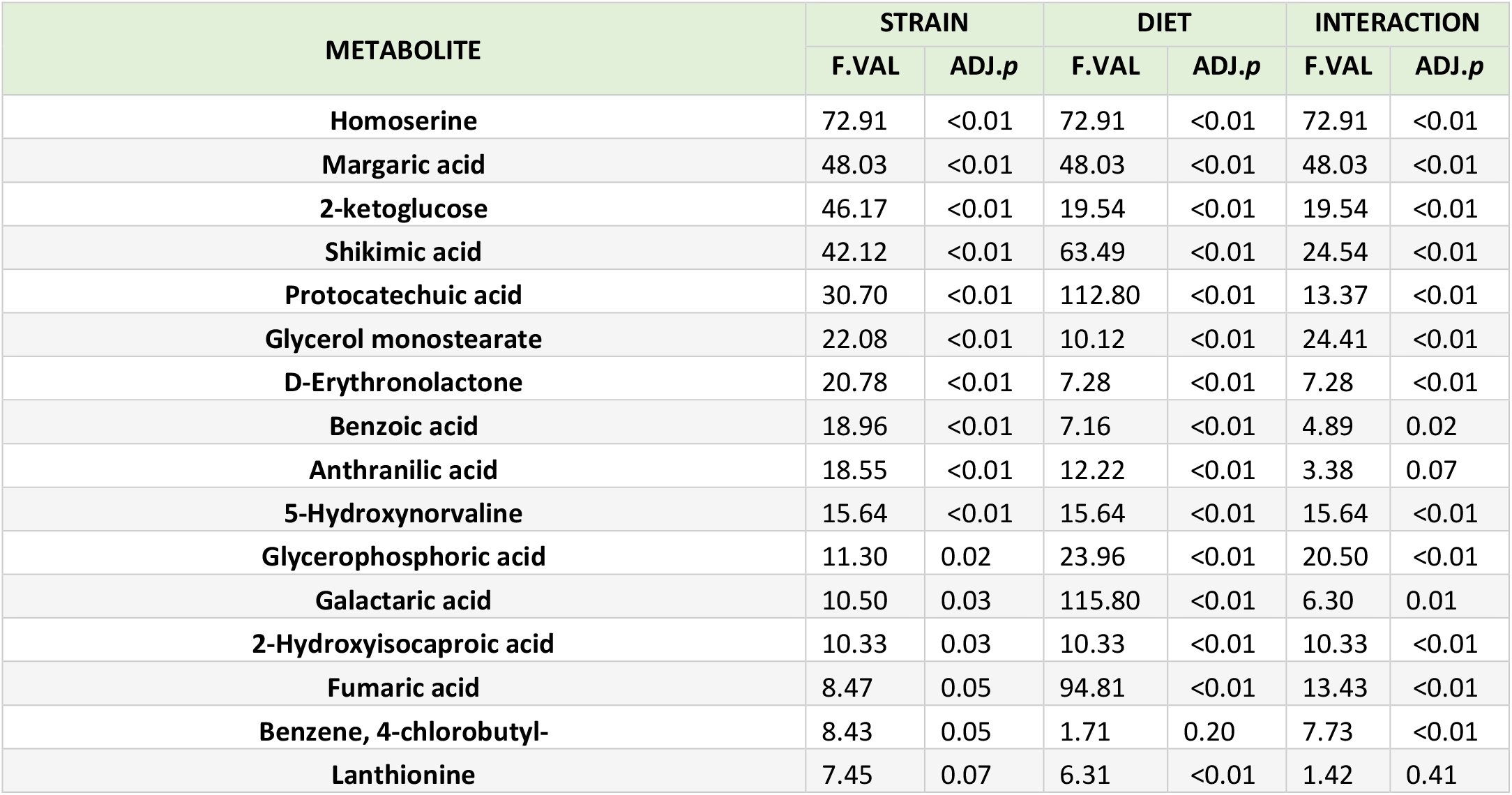
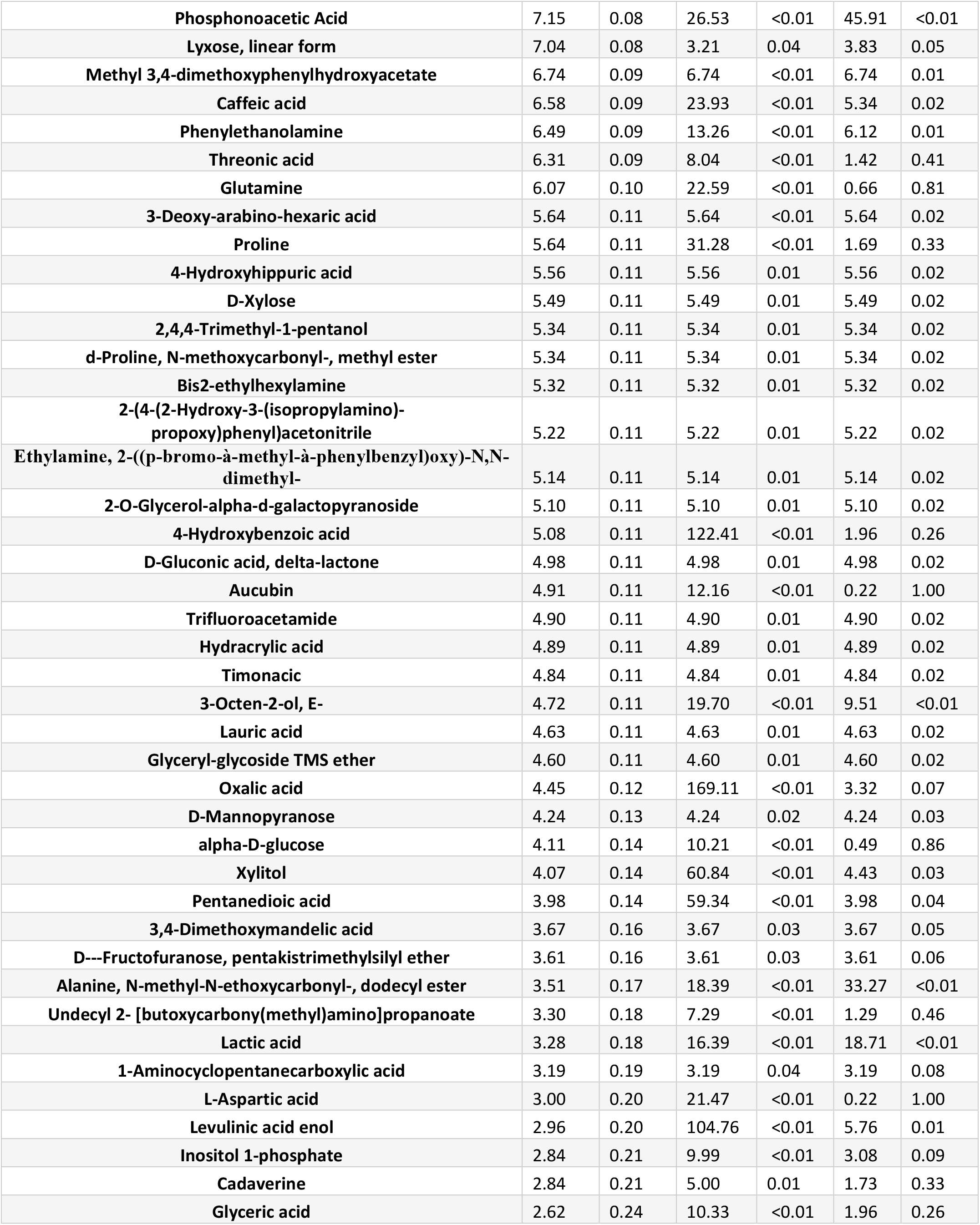
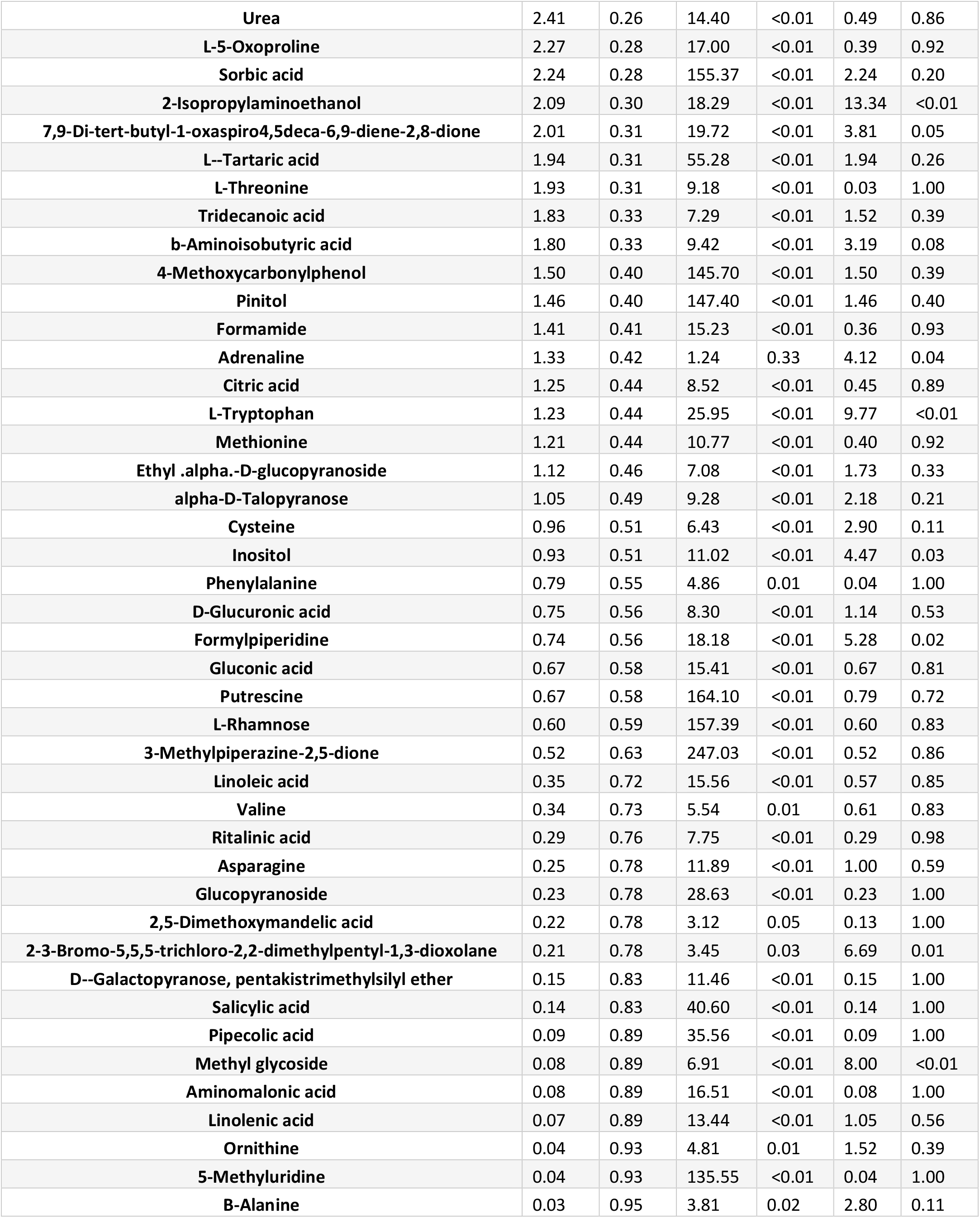
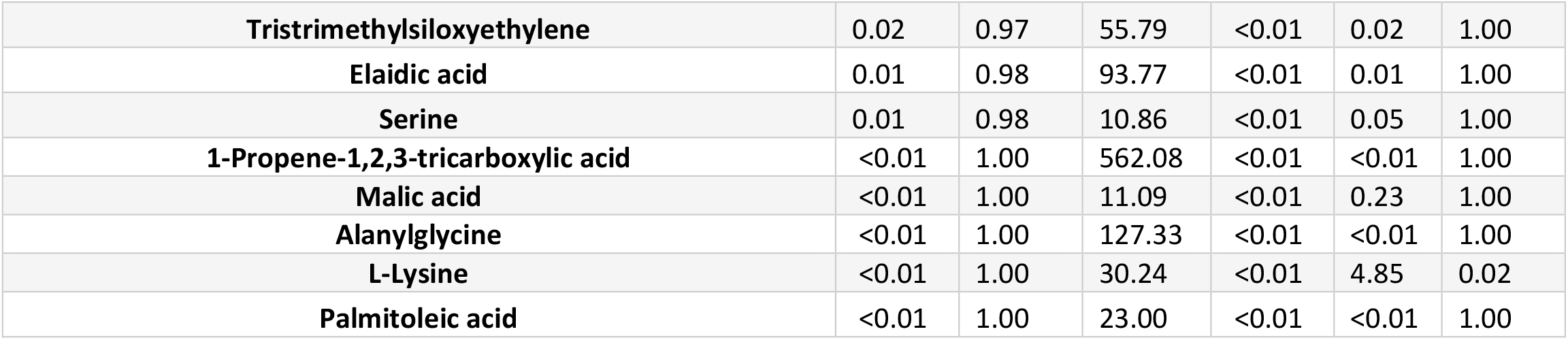
Significant features identified by two-way ANOVA. FDR’s correction was applied to adjust the p-values.

**Figure 2:**
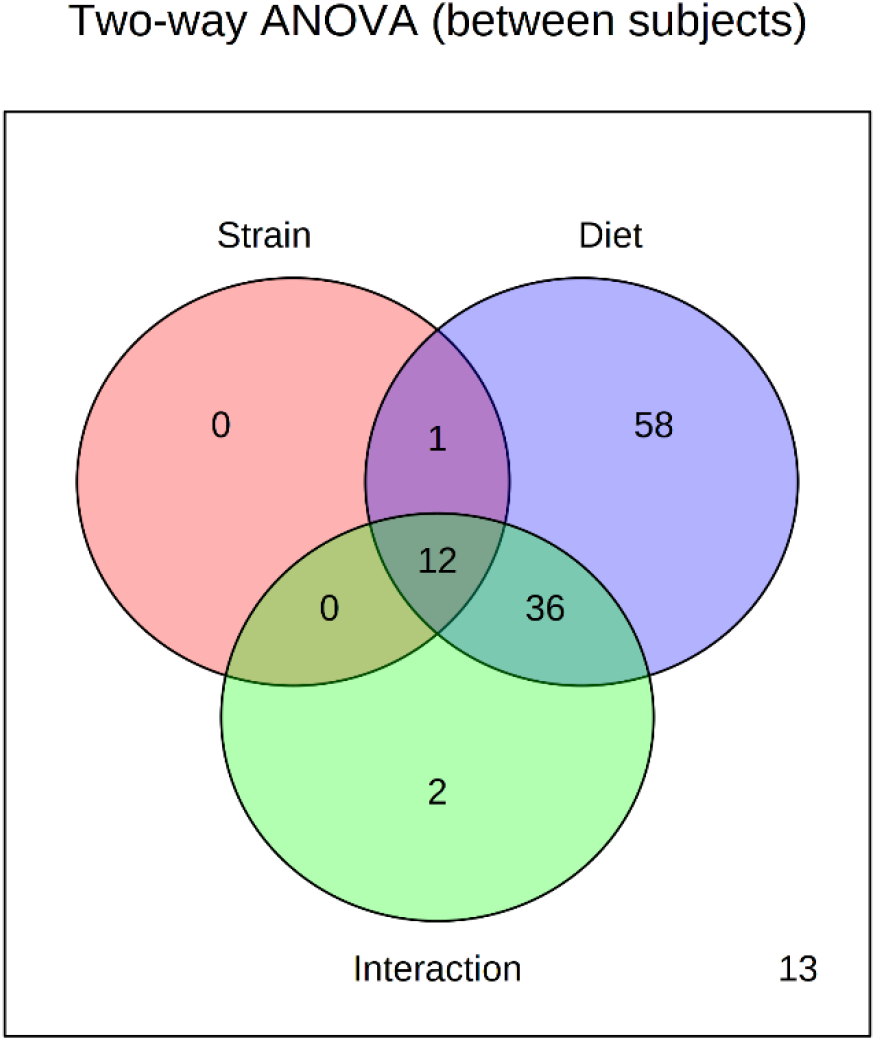
Venn diagram showing the important features of the midgut metabolome of *S. frugiperda* larvae selected by two-way ANOVA whose levels were affected by strain, diet or interaction of strain and diet.

In the heatmap, we also see two major clusters: one composed of metabolites mostly present in the gut of the larvae fed on the artificial diet (Fig. 3A) and another cluster composed of metabolites present in the larvae fed on the natural diets (Fig. 3B).

**Figure 3.**
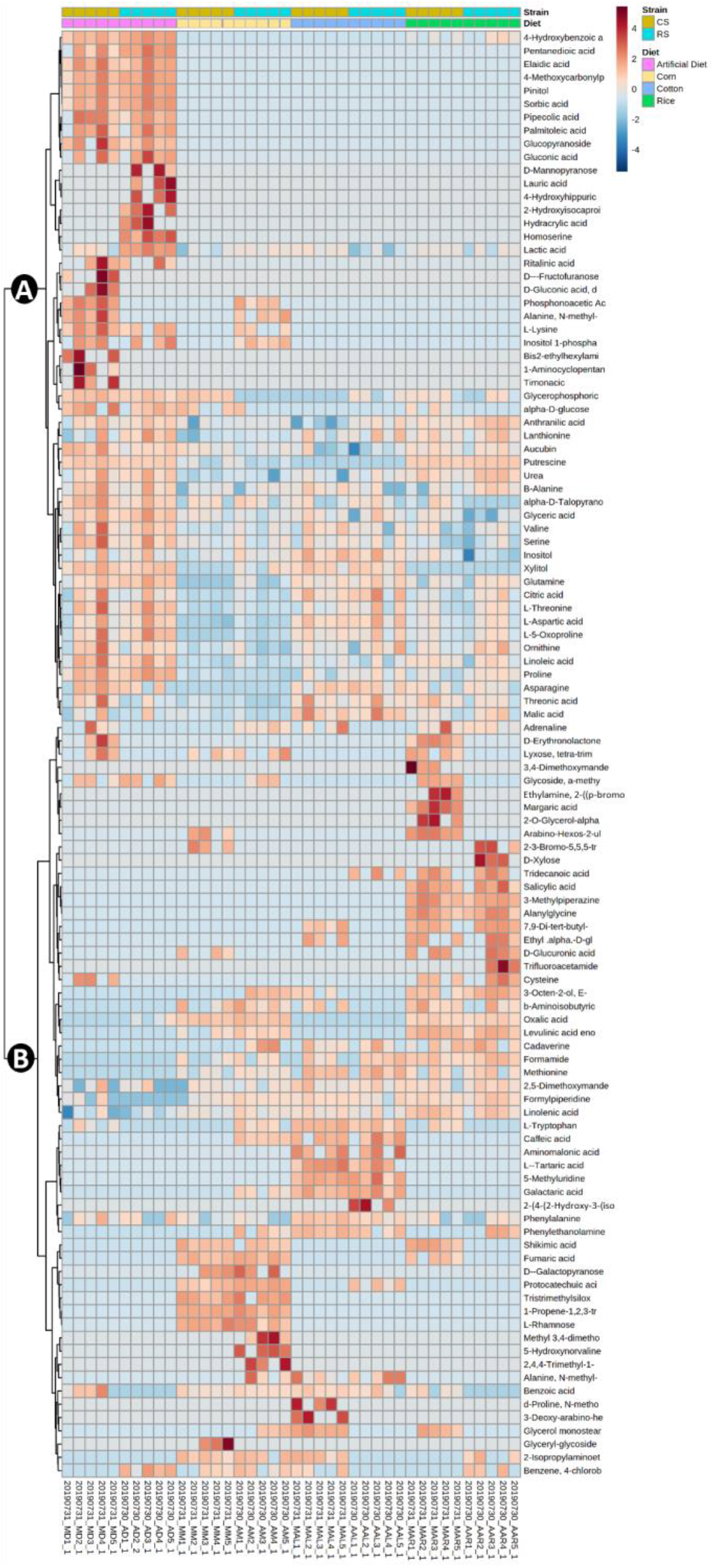
Heatmap of midgut metabolites from corn and rice strains of *Spodoptera frugiperda* larvae after feeding on artificial diet, corn, cotton, or rice. The experimental groups were rice strain on rice (RiRS), corn (CoRS), cotton (CtRS) and artificial diet (DiRS) and corn strain on the same diets (RiCS, CoCS, CtCS and DiCS).

Within the first cluster (Fig. 3A), there are four other sub clusters: one where the metabolites are similarly abundant in both strains, two sub clusters where the differences between strains are most evident, being one sub cluster composed of metabolites more abundant in *RS* and the second sub cluster composed of metabolites more abundant in *CS;* Finally, there is a sub cluster where besides the metabolites were abuntant in larvae fed on artificial diet, the metabolites were also abundant in the midgut of caterpillars fed on rice and cotton (Fig. 3A).

Whereas, in the second large cluster (Fig. 3B), we highlighted five sub-clusters: one in which the gut metabolomic profile was associated with *CS* when feeding on rice, another in which the profile was similar among the strains feeding on rice, the next was characterized by similar abundance among all plants, and the last two sub-clusters were characterized by metabolites associated with cotton and corn plants, respectively (Fig. 3B).

Pairwise analyses of the *RS* and *CS* gut metabolomes of larvae fed on each food source led to the identification of metabolites that differentiate FAW strains (Table 2). *RS* fed on corn had a higher abundance of caffeic acid, inositol 1-phosphate, an unidentified biogenic amine, protocatechuic acid, 5- hydroxynorvaline and 3-octen-2-ol, (E)- than the *CS* larvae. Only the abundance of glycerophosphoric acid was higher in *CS* than in *RS* larvae (Table 2). In cotton, the abundances of glycerol monostearate and 2-isopropylaminoethanol were higher in *CS* than in *RS* larvae. In rice-fed larvae, *CS* larvae had higher levels of shikimic acid, 2-ketoglucose, D-erythronolactone and margaric acid, while *RS* had higher anthranilic acid. The midgut of *RS* larvae fed on artificial diet were characterized by higher phosphonoacetic acid and alanine, N-methyl-N-ethoxycarbonyl-, dodecyl ester than in *RS*, while homoserine, lactic acid, and 4-chlorobutyl-benzene were more abundant in the midgut of *CS* larvae (Table 2).

**Table 2.**
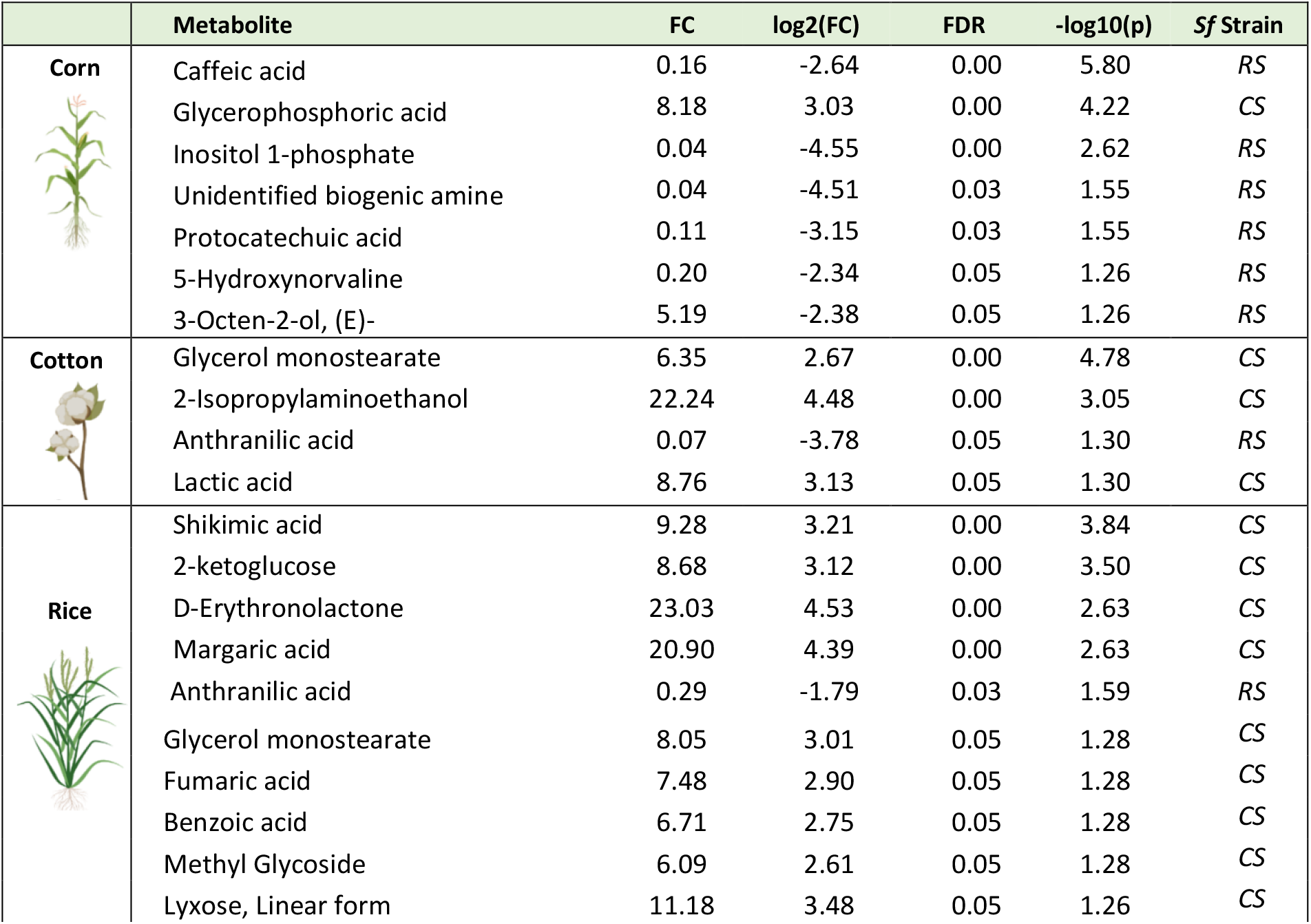
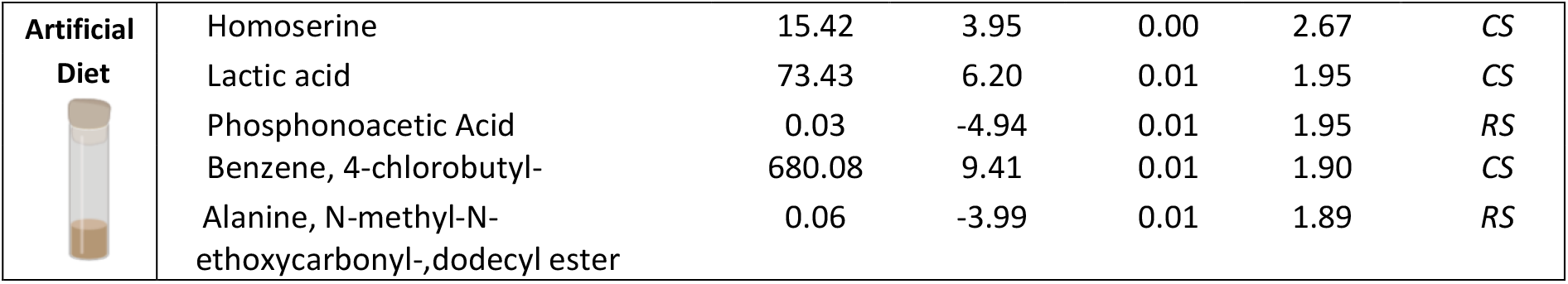
Significant features of midgut metabolome of *Spodoptera frugiperda* strains (*CS vs RS*) larvae after feeding on different food sources identified by Volcano plot with fold change threshold 2 and t-tests. False discovery rate correction was applied to adjust the p-values (p ≤ 0.05) and the strain where the compound were more abundant is identified.

|  | Metabolite | FC | log2(FC) | FDR | -log10(p) | Sf Strain |
| --- | --- | --- | --- | --- | --- | --- |
| **Corn** | Caffeic acid | 0.16 | -2.64 | 0.00 | 5.80 | *RS* |
|  | Glycerophosphoric acid | 8.18 | 3.03 | 0.00 | 4.22 | *CS* |
|  | Inositol 1-phosphate | 0.04 | -4.55 | 0.00 | 2.62 | *RS* |
|  | Unidentified biogenic amine | 0.04 | -4.51 | 0.03 | 1.55 | *RS* |
|  | Protocatechuic acid | 0.11 | -3.15 | 0.03 | 1.55 | *RS* |
|  | 5-Hydroxynorvaline | 0.20 | -2.34 | 0.05 | 1.26 | *RS* |
|  | 3-Octen-2-ol, (E)- | 5.19 | -2.38 | 0.05 | 1.26 | *RS* |
| **Cotton** | Glycerol monostearate | 6.35 | 2.67 | 0.00 | 4.78 | *CS* |
|  | 2-Isopropylaminoethanol | 22.24 | 4.48 | 0.00 | 3.05 | *CS* |
|  | Anthranilic acid | 0.07 | -3.78 | 0.05 | 1.30 | *RS* |
|  | Lactic acid | 8.76 | 3.13 | 0.05 | 1.30 | *CS* |
| **Rice** | Shikimic acid | 9.28 | 3.21 | 0.00 | 3.84 | *CS* |
|  | 2-ketoglucose | 8.68 | 3.12 | 0.00 | 3.50 | *CS* |
|  | D-Erythronolactone | 23.03 | 4.53 | 0.00 | 2.63 | *CS* |
|  | Margaric acid | 20.90 | 4.39 | 0.00 | 2.63 | *CS* |
|  | Anthranilic acid | 0.29 | -1.79 | 0.03 | 1.59 | *RS* |
|  | Glycerol monostearate | 8.05 | 3.01 | 0.05 | 1.28 | *CS* |
|  | Fumaric acid | 7.48 | 2.90 | 0.05 | 1.28 | *CS* |
|  | Benzoic acid | 6.71 | 2.75 | 0.05 | 1.28 | *CS* |
|  | Methyl Glycoside | 6.09 | 2.61 | 0.05 | 1.28 | *CS* |
|  | Lyxose, Linear form | 11.18 | 3.48 | 0.05 | 1.26 | *CS* |
| **Artificial Diet** | Homoserine | 15.42 | 3.95 | 0.00 | 2.67 | CS |
|  | Lactic acid | 73.43 | 6.20 | 0.01 | 1.95 | CS |
|  | Phosphonoacetic Acid | 0.03 | -4.94 | 0.01 | 1.95 | RS |
|  | Benzene, 4-chlorobutyl- | 680.08 | 9.41 | 0.01 | 1.90 | CS |
|  | Alanine, N-methyl-N-ethoxycarbonyl-,dodecyl ester | 0.06 | -3.99 | 0.01 | 1.89 | RS |

## 4. DISCUSSION

The metabolic profile of the FAW larvae midgut is largely influenced by the food source used, and the two strains differ in every food source analyzed. Our data demonstrates the *RS* and *CS* interact differently with the substrate on which they are feeding, potentially due to differential metabolism of plant chemistry (Silva-Brandão et al., 2017). Differences at the genomic level are reported for these strains (Dumas et al., 2015), particularly with the large variation they have in the number of copies of genes and gene sequences encoding for detoxification and digestive enzymes (Gouin et al., 2017). In addition, our results are also consistent with the plethora of differences at the transcriptional level reported for the whole body of both strains when feeding on the same host plants. These strains were demonstrated to have differences in the expression levels of genes encoding for proteins with oxidoreductase activity, metal-ion binding, and hydrolase activity, which are also related to the metabolism of xenobiotics (Orsucci et al., 2020; Silva-Brandão et al., 2017).

Some compounds reported here in the FAW midgut have a defensive function in plants against insects, such as shikimic acid. This compound has been shown to reduce intestinal proteolytic activity in insects by acidifying the intestinal lumen. However, some specialist insects such as *Gilpinia hercyniae* (Hymenoptera) are able to metabolize and neutralize the effect of this compound, through their gut bacteria (Schopf, 1986). Therefore, the higher abundance of shikimic acid in the *CS* larval midgut comparing to *RS* feeding on rice indicates the *CS* has a lower capacity to process this metabolite. Additionally, the difference in shikimic acid metabolization may also be due to the differential activity of the gut microbiota of the strains, similar to the adaptation found in *G. hercyniae* (Jensen, 1991). Likewise, margaric acid, or heptadecanoic acid, was shown to accumulate in *CS* when feeding on rice. This compound was negatively correlated with oviposition, eclosion, and nymphal survival of *Stephanitis pyrioides* (Hemiptera: Tingidae) on azalea (*Rhododendron sp*.), but positively correlated with duration of development, indicating an arrestment of the developmental period (Wang et al., 1999). Margaric acid is also found in rice plants (Jones et al. 2011) so it may play a role in the low performance and survival of *CS* on rice plants (Silva-Brandão et al., 2017).

The reverse pattern was observed for strains feeding on corn, where the defensive metabolites, 5-hydroxynorvaline, caffeic acid and protocatechuic acid, where measured in higher abundance in *RS* larval midgut than in *CS*. The accumulation of 5-hydroxynorvaline in maize leaves has been demonstrated after the feeding of *S. exigua* and the aphid, *Rhopalosiphum maidi* (Yan et al., 2015), suggesting that this metabolite can provide protection against herbivores. When this compound was added to the artificial diet, it reduced aphid growth and reproduction, but no significant effect was found on *S. exigua* larval growth. However, 5-hydroxynorvaline also plays a defensive role by replacing amino acids in protein synthesis or by inhibiting the biosynthetic pathways of many microorganisms(Guirard, 1958; Heremans and Jacobs, 1994; Huang et al., 2011; Kurtin et al., 1971; Washtien et al., 1977). Thus, it is possible 5-hydroxynorvaline negatively impact the gut bacteria and impairs its contribution to the host.

The flavonoids caffeic and protocatechuic acids are referred as potential insecticides due to their toxic effects (War et al., 2013). *Helicoverpa armigera* larvae fed on caffeic and protocatechuic acids displayed reduced digestive and detoxification activity due to a reduction in serine protease, trypsin, and esterase activity. The larvae also showed greater reduction in larval weight and higher mortality when compared to the larvae fed on untreated control diet(War et al., 2013). Moreover, caffeic acid also increases the oxidative stress in the gut of insect herbivores due to the elevation of protein oxidation, lipid peroxidation products and release of free ions(Summers and Felton, 1994). The accumulation of these flavanoids in the *RS* larval midgut could explain why *RS* does not perform as well as the *CS* when feeding on maize (Orsucci et al., 2020; Silva-Brandão et al., 2017).

The lower levels of defensive plant compounds in the midgut of the strains when they were feeding on their preferred host plants (*RS* on and rice *CS* on maize) and their higher abundancies in the midgut when larvae were feeding on the non-preferred host plants (Fig. S1), suggest either differential metabolization of the food source as discussed above and/or differential elicitation of metabolic response in the host plant. Additionally, FAW strains were demonstrated inducing different defense responses in maize and Bermuda grass via specific differences in their saliva composition (Acevedo et al., 2018). The gut-associated microbes in their oral secretions also play a role mediating the insect-plant interaction by regulating plant defenses upon their secretion through insect oral secretions (Acevedo et al., 2017).

Our findings also suggest that the strains of FAW metabolize the artificial diet differently. The diet has been widely used for several Lepidoptera, including FAW, demonstrating good performance (Gardner et al., 1984; Perkins, 1979; Silva et al., 2018), However, most experiments were performed using the corn strain. There is only one study as far as we know showing that *CS* larvae were significantly heavier than *RS* larvae when they were fed the artificial diet (Silva-Brandão et al., 2017). Furthermore, artificial diets generally provide unrealistic amounts of soluble carbohydrates, proteins, and fats. Perhaps the reason for a greater accumulation of compounds in the larval gut when compared to other diets is due to the large amount of these compounds in the food, which does not allow their complete metabolization.

The fact that only glycerol monostearate and 2-isopropylaminoethanol were differentially abundant between *CS* and *RS* strains, being both more abundant in the midgut of *CS* larvae suggest that the strains behave in a very similar way when feeding on this plant. Interesting, it is suggested that feeding on dicot is a primitive condition of the FAW complex and feeding on grasses is a more recent event (Kergoat et al., 2012). Additionally, studies also demonstrated that FAW presents low performance and low survival rate when feeding on cotton (Ali et al., 1990; Barros et al., 2010)

In conclusion, our study documented the effects of host strains and dietary on the metabolome of the FAW midgut. Our analyses found not only diet effects on the metabolome but indicate differential digestive metabolism between FAW strains. and identified marker metabolites that may help us to better understand the mechanisms involved in host adaptation. Our results shed light on our understanding of metabolic activities in the FAW, being a unit composed of its own metabolome and the metabolome of the associated gut microbiota. Further analyses are essential to reveal the links between gut microbiota composition and host metabolic phenotype, thus providing a holistic understanding of the functionality and adaptability of strains to host plants.

## 5. ACKNOWLEDGEMENTS

We are grateful to the technician at the Insect Interactions Laboratory, Marcele Coelho for her help in the rearing of *S. frugiperda* and the host plant management. We also thank the technicians of the multi-User Proteomics, Metabolomics and Lipidomics laboratory, Thais Cataldi and Monica Labate for their help with the gas chromatography and mass spectrometry analyses.

## 9. SUPPLEMENTARY MATERIAL

**Figure S1.**
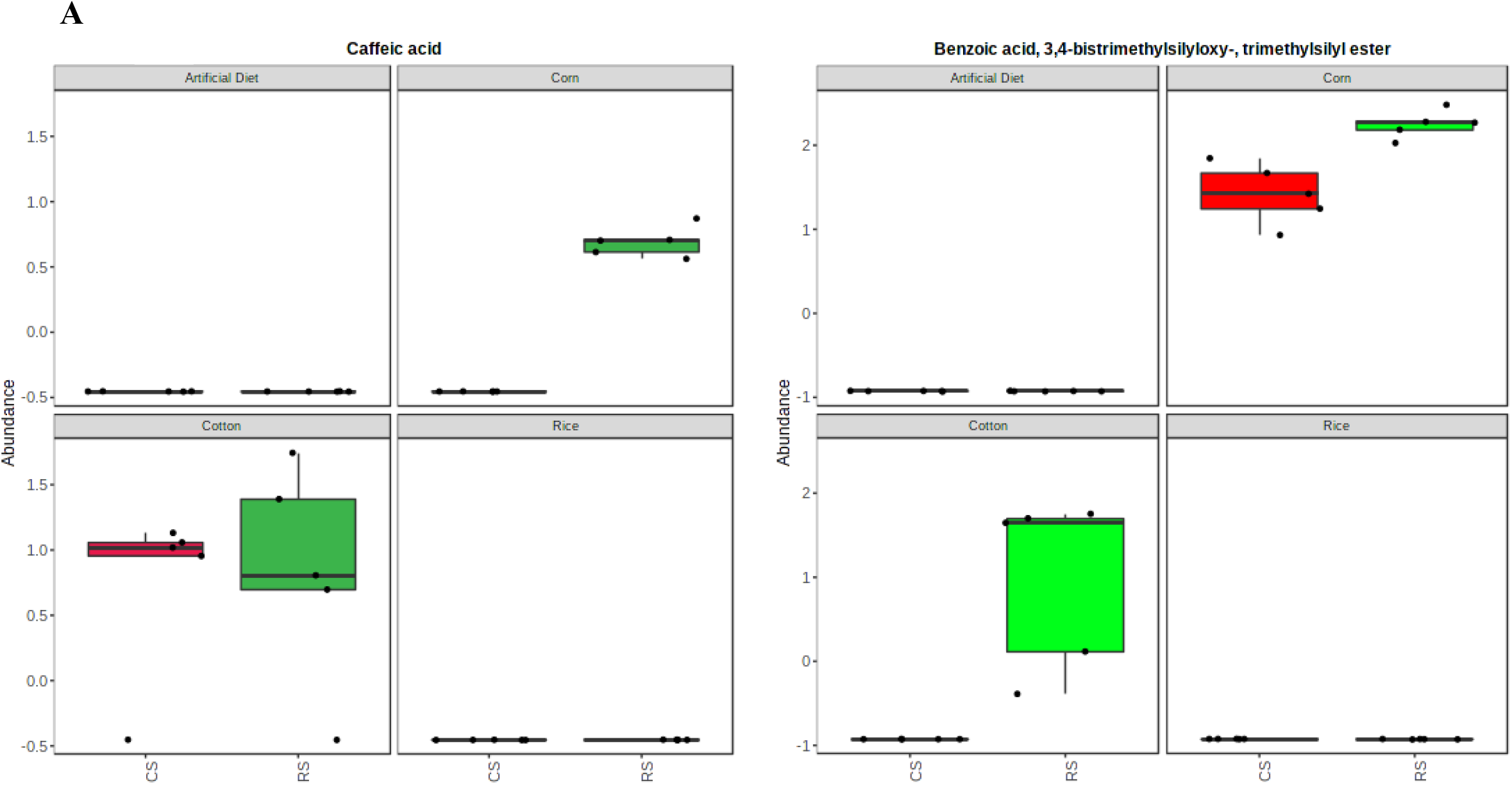
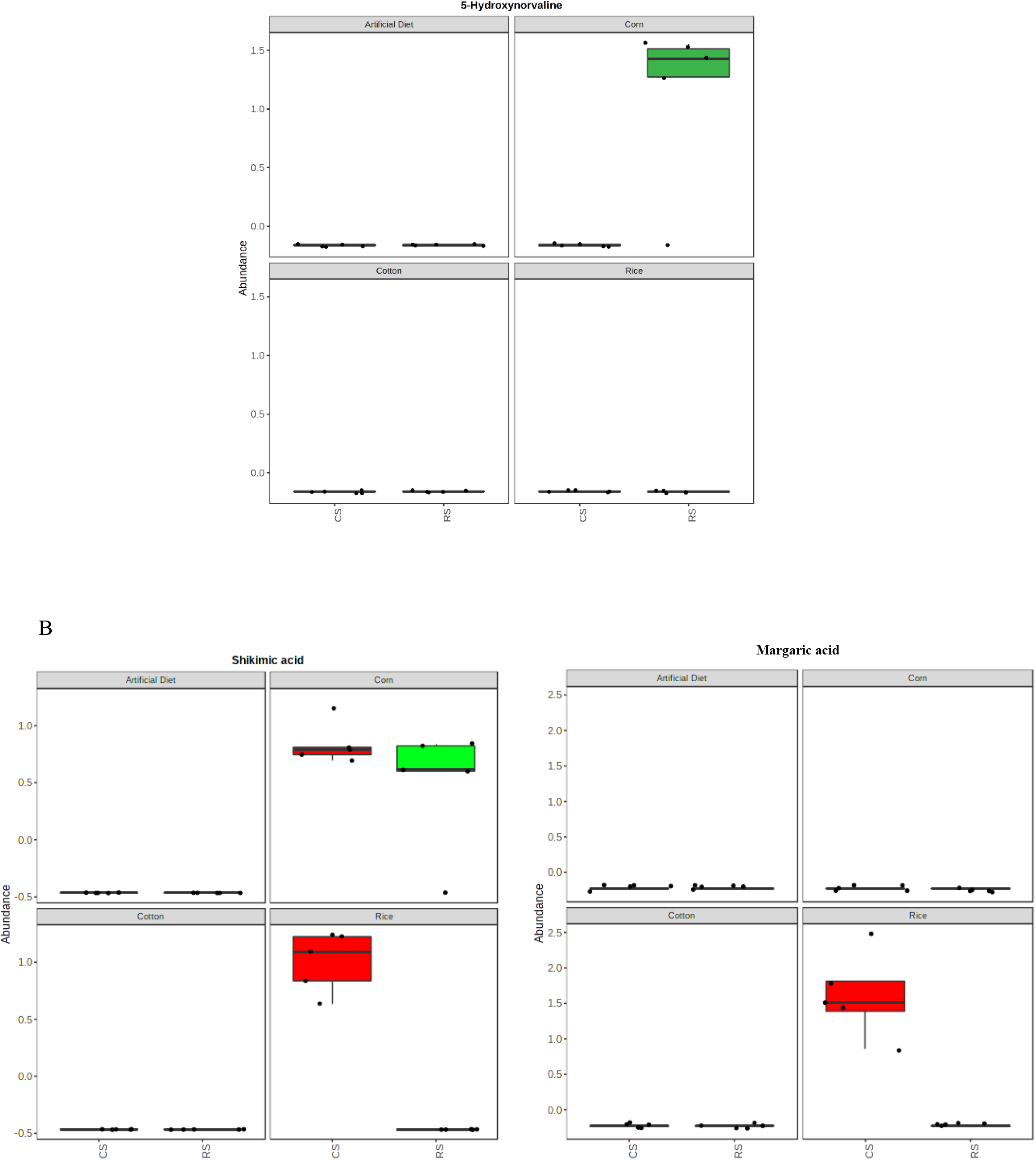
Boxplot of significant features of midgut metabolome of *Spodoptera frugiperda* strains larvae after feeding on maize (A) and rice (B) identified by Volcano plot with fold change threshold 2 and t-tests threshold 0.05. False discovery rate correction was applied to adjust the p-values (0.05).

